# A protective role for B-1 cells and oxidation-specific epitope IgM in lung fibrosis

**DOI:** 10.1101/2024.04.11.589137

**Authors:** Jeffrey M. Sturek, Riley T. Hannan, Aditi Upadhye, Eva Otoupalova, Edwin T. Faron, Amr A.E. Atya, Cassidy Thomas, Vernerdean Johnson, Andrew Miller, James C. Garmey, Marie D. Burdick, Thomas H. Barker, Alexandra Kadl, Yun M. Shim, Coleen A. McNamara

**Affiliations:** Division of Pulmonary & Critical Care Medicine, Department of Medicine, University of Virginia School of Medicine; Cardiovascular Research Center, University of Virginia School of Medicine; Beirne B. Carter Center for Immunology Research, University of Virginia School of Medicine; University of Virginia School of Medicine; Department of Biomedical Engineering, University of Virginia School of Medicine

## Abstract

Idiopathic pulmonary fibrosis (IPF) is a morbid fibrotic lung disease with limited treatment options. The pathophysiology of IPF remains poorly understood, and elucidation of the cellular and molecular mechanisms of IPF pathogenesis is key to the development of new therapeutics. B-1 cells are an innate B cell population which play an important role linking innate and adaptive immunity. B-1 cells spontaneously secrete natural IgM and prevent inflammation in several disease states. One class of these IgM recognize oxidation-specific epitopes (OSE), which have been shown to be generated in lung injury and to promote fibrosis. A main B-1 cell reservoir is the pleural space, adjacent to the typical distribution of fibrosis in IPF. In this study, we demonstrate that B-1 cells are recruited to the lung during injury where they secrete IgM to OSE (IgM^OSE^). We also show that the pleural B-1 cell reservoir responds to lung injury through regulation of the chemokine receptor CXCR4. Mechanistically we show that the transcription factor *Id3* is a novel negative regulator of CXCR4 expression. Using mice with B-cell specific Id3 deficiency, a model of increased B-1b cells, we demonstrate decreased bleomycin-induced fibrosis compared to littermate controls. Furthermore, we show that mice deficient in secretory IgM (*sIgM^-/-^*) have higher mortality in response to bleomycin-induced lung injury, which is partially mitigated through airway delivery of the IgM^OSE^ E06. Additionally, we provide insight into potential mechanisms of IgM in attenuation of fibrosis through RNA sequencing and pathway analysis, highlighting complement activation and extracellular matrix deposition as key differentially regulated pathways.

## INTRODUCTION

Idiopathic pulmonary fibrosis (IPF) is a progressive fibrotic lung disease with a median survival of three years, a rate worse than many malignancies^1, 2^. Recently medical therapies have become available which slow disease progression^3, 4^, but the only definitive therapy remains lung transplantation. The pathogenesis of IPF has been enigmatic, limiting the development of targeted therapeutics. The current working hypothesis is that IPF results from an aberrant tissue repair response to repetitive micro-injury (identified or unidentified) in the context of genetic susceptibility with involvement of both innate and adaptive immune mechanisms^5^. Broad immunosuppression, however, has been shown to be deleterious in IPF^6^. Therefore, understanding the immunologic response to lung injury and how this impacts the development of fibrosis may provide key insights into IPF pathogenesis and reveal avenues for novel therapeutic modalities.

B-1 cells are an innate B cell population which produce the majority of the circulating IgM in a non-infected state^7^. B-1 cells can be sub-classified based on CD5 expression, with B-1a cells being CD5 positive, and B-1b CD5 negative. The immunoglobulins secreted by B-1 cells are referred to as natural antibodies (NAbs) as their production is spontaneous and independent of T cell help. They are evolutionarily conserved and recognize exogenous (pathogen-associated molecular patterns or PAMPs) as well as modified endogenous (damage-associated molecular patterns or DAMPs) epitopes, such as oxidized lipids, apoptotic cells, and leukocytes^8, 9^. One such class of epitopes are referred to as oxidation-specific epitopes (OSE). These are DAMPs formed through phospholipid oxidation and exposed on the surface of apoptotic cells in the setting of inflammation. Unlike high-specificity, high-affinity IgG, IgM^OSE^ and other NAbs are relatively low-affinity and can recognize multiple different epitopes. They are considered to be protective both in terms of providing an early innate response to infection, and by masking autoantigens and preventing autoimmunity and inflammation^8, 10^. B-1 cells and IgM^OSE^ have been shown to play a protective role in a number of inflammatory disease states including chronic diseases such as atherosclerosis^9, 11^, obesity^12^, and systemic lupus erythematosis^13^; and in acute inflammation such as ischemia reperfusion injury^14^. B-2 cells on the other hand, are considered pro-inflammatory^15^. B-1 cells and NAbs have also been shown to play a role in response to various pulmonary infections^16, 17^, and sepsis-induced acute lung injury^18^ however little is known about their role in non-infectious pulmonary inflammation or fibrosis.

The lung is an ideal site for an evolutionarily conserved role of IgM^OSE^. The highly oxidative environment of the lung has been shown to produce higher levels of OSEs, such as oxidized phospholipids, both in the normal physiological state as well as in increased inflammation and oxidative stress^19, 20^. Additionally, in the mouse the main reservoir of B-1 cells is serosal cavities, including the pleural cavity^8^, where they are poised to respond to pulmonary antigens and migrate to sites of infection or injury. This is of particular interest as the distribution of fibrosis in IPF is peripheral and basilar, immediately adjacent to the B-1 cell niche.

Recent genetics studies suggest that host defense and innate immune signaling through toll-like receptors (TLRs) may play a key role in IPF. TLRs are a group of transmembrane proteins present on the surface of immune cells which bind to a range of PAMPs and DAMPs and thereby activate an immune response. Genetic mutations in TLR signaling have been shown to be associated with IPF susceptibility and survival^21^. What TLR ligands may play a role in IPF pathogenesis is unknown, though there may be multiple, and the response is likely complex. OSEs are a particularly interesting candidate group. They have been extensively studied in the context of atherosclerosis where they activate the innate immune system and promote atherogenesis^22^. Importantly, OSEs have been shown to be formed in lung injury and inflammation and to promote fibrosis^23–25^. To-date this work has largely focused on the role of TLRs and scavenger receptors (SRs) in this process.

Finally, like in atherosclerosis, in addition to these innate immune mechanisms, there is also evidence for a role for B cells in IPF. Importantly, none of these studies to-date have differentiated between the effects of B-1 versus B-2 cells and based on their various approaches almost universally represent primarily B-2 cells. The results of studies of the effect of B cells on lung fibrosis is mixed, depending on the mode of B cell modulation. Some murine studies have shown that B cells are recruited to the lung during injury and that this promotes fibrosis^26^, while other studies have demonstrated a protective role for B cells^27^ or no effect^28^.

Surgical lung biopsy specimens of patients with IPF have been shown to have increased numbers of B cells which accumulate as the disease progresses, as well as increased staining of IgG^29–31^. Furthermore, circulating levels of the B cell survival and differentiation factor BlyS (also known as BAFF or TNFSF13B) have been shown to be increased in IPF and were associated with increased mortality^29^.

In summary, there is ample evidence for the role of B-cells and oxidation-specific epitope responses in pathogenesis of IPF. However, the exact link between products of oxidative injury and adaptive immune responses in pathogenesis of lung fibrosis is unknown. In the current study, we examine the role of B-1 cells and IgM^OSE^ in lung injury and fibrosis.

## RESULTS

### B-1 cells are recruited to the lung during bleomycin-induced lung injury where they secrete natural IgM

To determine the role of B-1 cells in the response to lung injury and fibrosis, we utilized the well-established bleomycin model^32^. The mechanism of action involves DNA damage through generation of superoxide and hydroxide free radicles. This has been shown to produce a robust pro-inflammatory response with production of a number of pro-inflammatory cytokines (such as IL-1, TNF-α, and IL-6) followed by an increase in pro-fibrotic markers (such as TGF-β1 fibronectin, and pro-collagen). Flow cytometry of bronchoalveolar lavage (BAL) and whole lung single cell suspensions (Supplemental Figure 1, gating strategy) showed that B-1 cells are recruited to the lung and airspace in bleomycin-induced lung injury (Figure 1). This was accompanied by increased levels of total IgM as well as for two representative IgM^OSE^, E06 and MDA-LDL in the BAL as measured by ELISA (Figure 2). To corroborate these findings and to test for functional IgM secretion, we performed enzyme-linked immunospot (ELISPOT) assays on single-cell suspensions from bronchoalveolar lavage (BAL), lung, and bone marrow similar to previously published^11^. These studies showed that bleomycin increased the number of IgM-secreting cells specifically in the lung, with no significant effect on the bone marrow, another site of secretory B-1 cells (Figure 2E-G). This was accompanied by increased mRNA levels of E06 in the lung (Figure 2H).

**Figure 1.**
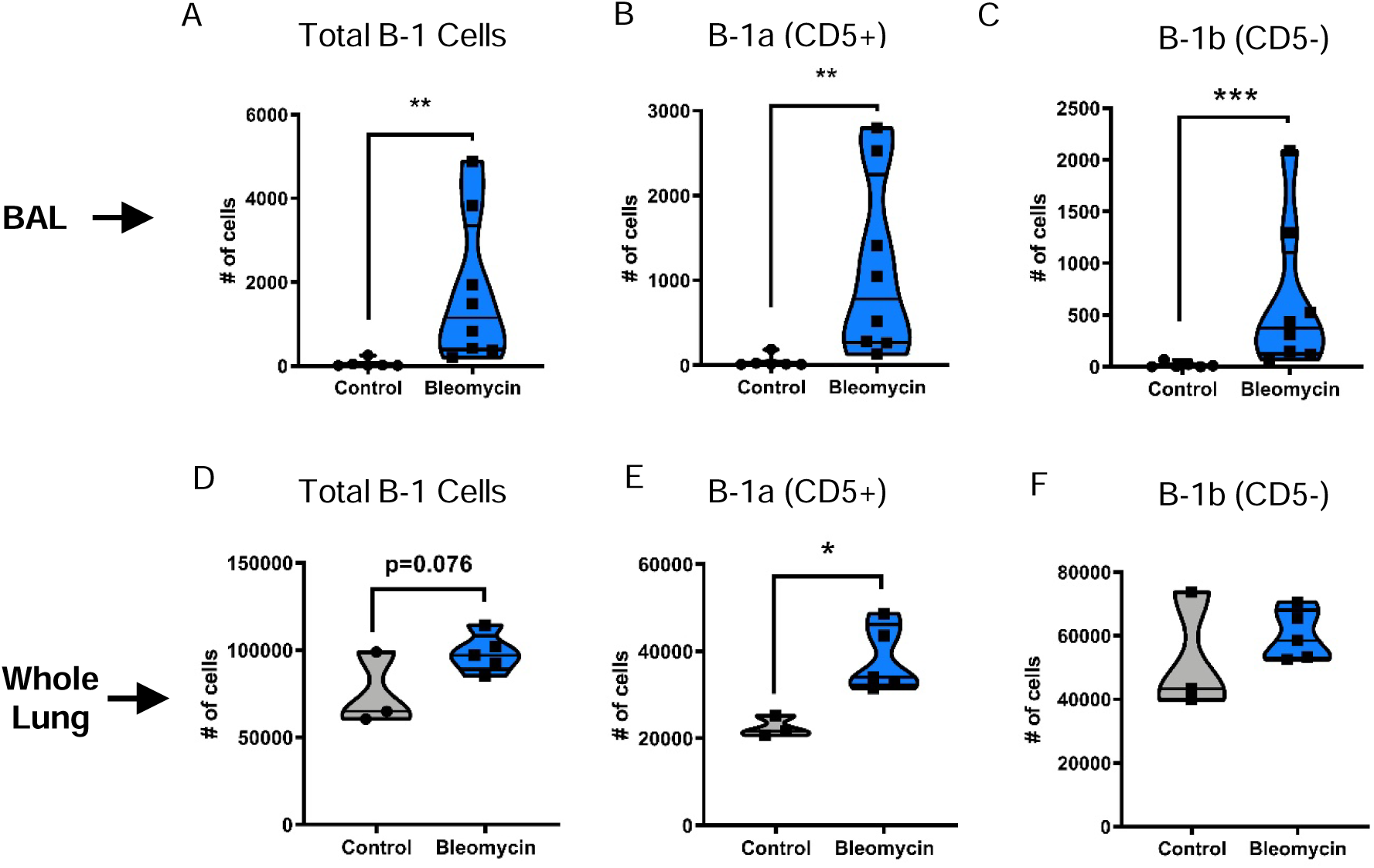
B-1 cells are recruited to the lung during bleomycin-induced lung injury. Mice were treated with bleomycin and bronchoalveolar lavage (BAL) and whole lung tissue were isolated 7 days following lung injury. Lung tissue was digested to single-cell suspension. B-1 cell numbers were quantified by flow cytometry in the BAL (A-C) and whole lung digest (D-F). n=3-8 per group, replicated over at least 2 separate experiments. *=p<0.05; **=p<0.01; ***=p<0.001

**Figure 2.**
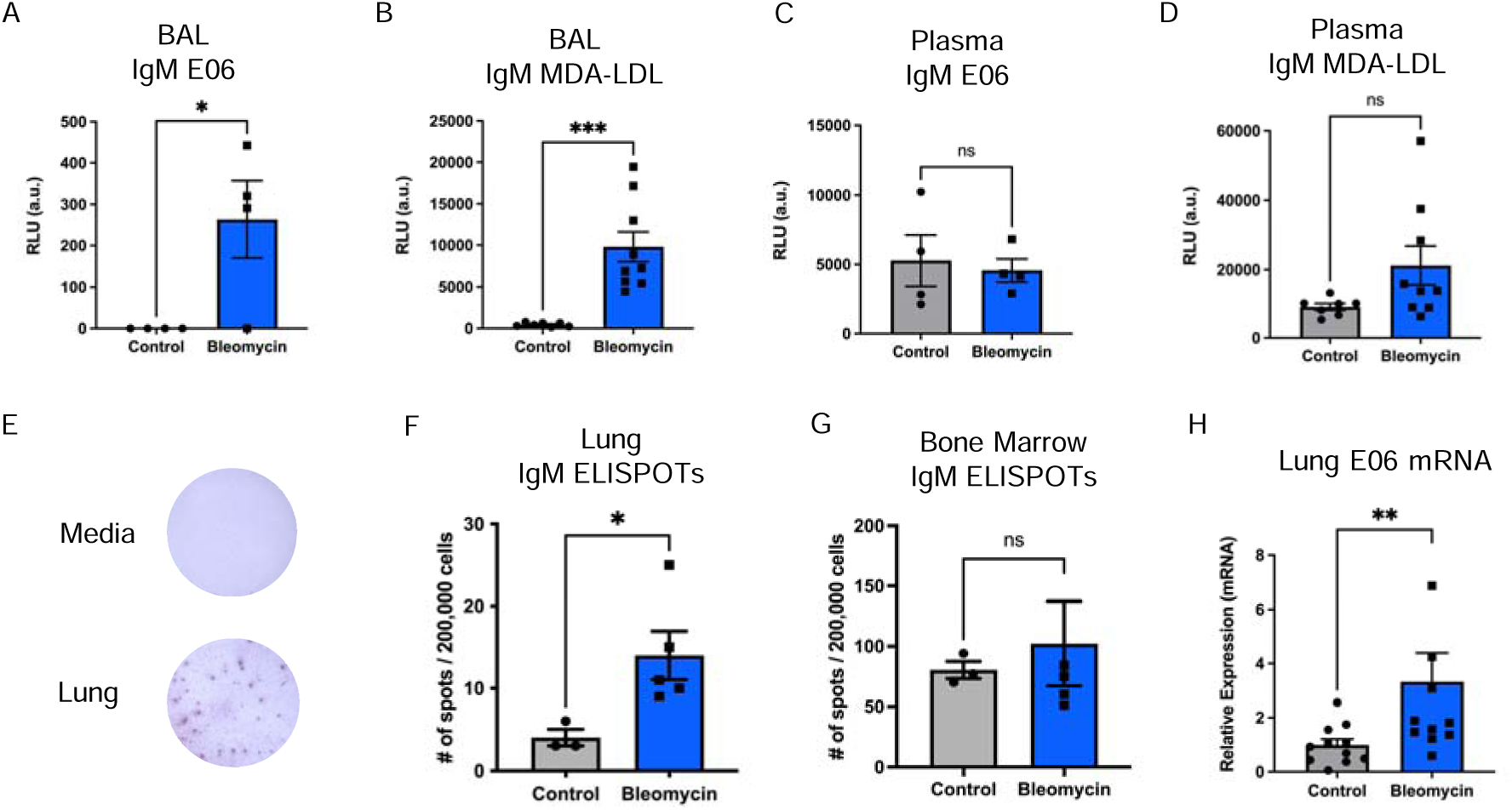
Bleomycin-induced lung injury promotes both total and oxidation-specific epitope IgM secretion specifically in the lung. ELISAs for IgM E06 and IgM MDA-LDL were performed on BAL (A,B) and plasma (C,D) from control and bleomycin-treated mice. E) Representative images of ELISPOT wells. IgM ELISPOT quantification from lung (F) and bone marrow (G) at 7 days following bleomycin. H) RT-PCR of lung tissue for E06 mRNA. n=3-9 per group. *=p<0.05; **=p<0.01; ***=p<0.001

To determine what factors might contribute to B-1 cell homing to the lung in bleomycin-induced lung injury, we measured surface expression of the chemokine receptor (CR) C-X-C chemokine receptor type 4 (CXCR4) in bleomycin-treated mice. Prior work from our laboratory has shown that CXCR4 is a key CR regulating B-1 cell homing^33^. Importantly, other prior published studies have also shown that the ligand for CXCR4, C-X-C motif chemokine ligand 12 (CXCL12) is increased in the injured lung and promotes leukocyte infiltration via CXCR4^34, 35^.

Data from these experiments show that CXCR4 is upregulated, both on lung B-1 cells as well as resident pleural B-1 cells, each approaching levels seen on infiltrating BAL B-1 cells (Figure 3). This suggests that the pleural space may provide a reservoir of B-1 cells which respond to chemotactic signals and migrate to the site of injury.

**Figure 3.**
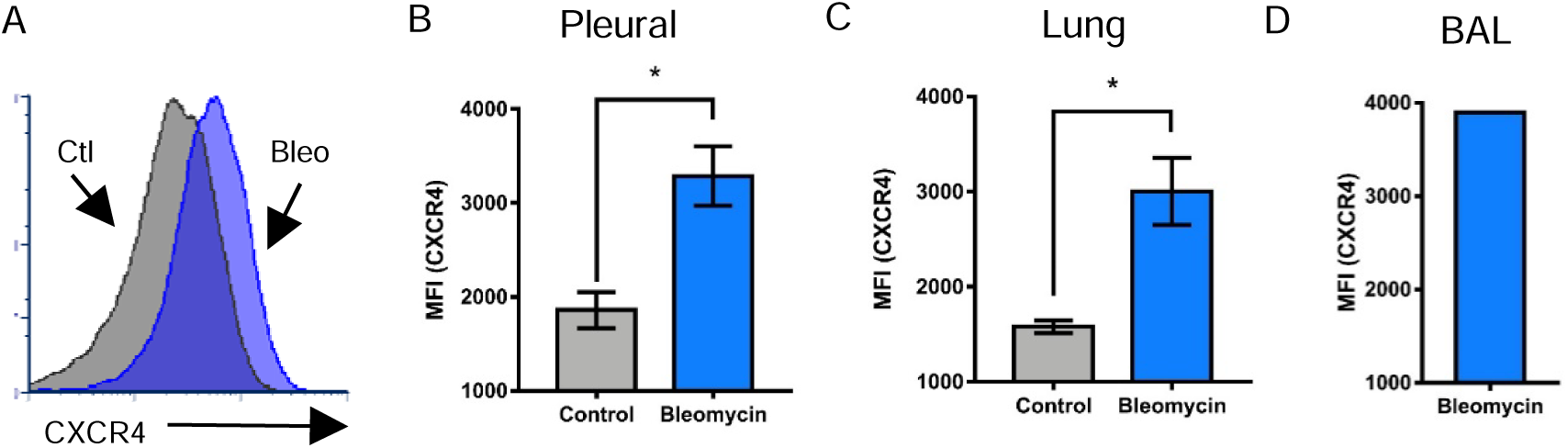
Upregulation of surface CXCR4 expression on pleural and lung B-1 cells. Surface expression of CXCR4 on B-1 cells as measured by flow cytometry. (A) representative histograms of CXCR4 expression on pleural B-1 cells. Median fluorescent intensity (MFI) of CXCR4 on pleural (B), lung (C), and BAL (D) B-1 cells. Note, insufficient numbers of B-1 cells are present in the BAL of control mice to permit quantification. n=3-5, replicated over two separate experiments. *=p<0.05.

### B cell-specific Id3 deficiency is a model of augmented pulmonary B-1 cells and natural IgM

To determine what effect B-1 cells and IgM^OSE^ may have on pulmonary fibrosis, we turned to mice deficient in the protein Inhibitor of Differentiation 3 (*Id3^-/-^*). Id3 is a member of the basic helix-loop-helix family of transcriptional regulators. Prior work in our laboratory has shown that Id3 regulates B-1b cell number, and that loss of Id3 function, in both mice and humans, leads to increased numbers of B-1 cells and IgM^OSE^ ^11, 12^. To test whether this also applied to the lung, we comprehensively evaluated B-1b cell distributions in *Id3^-/-^* mice over the course of bleomycin-induced lung injury. These studies confirmed that, like in other compartments, loss of Id3 leads to increased numbers of B-1b cells and increased IgM^OSE^ in the lung (Figure 4), both in global as well as B cell-specific Id3 deficiency, henceforth referred to as Id3^BKO^ (*Id3^flox/flox^;CD19^cre/+^*).

**Figure 4.**
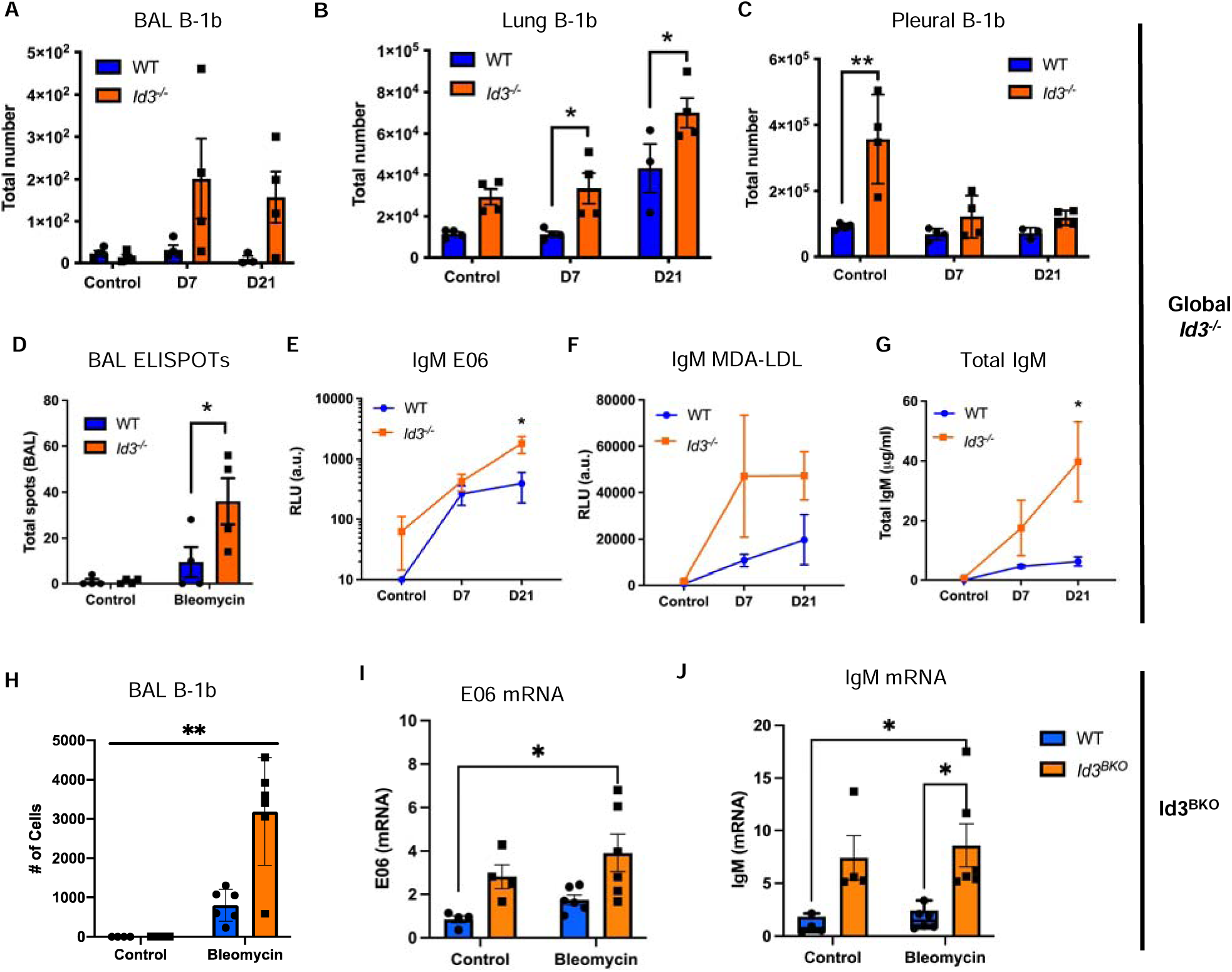
Id3-deficiency is a model of increased pulmonary B-1b cells and IgM^OSE^. *Id3^-/-^* mice and wild-type (WT) littermate controls were treated with bleomycin. BAL, lung, and pleural lavage were harvested at day 0, 7, or 21. Flow cytometry for B-1b cells (A-C) and ELISPOTs for IgM secreting cells (D) were performed. The BAL supernatant was also assayed for total IgM and IgM^OSE^ by ELISA (E-G). *Id3^BKO^*mice were treated with bleomycin and harvested at day 21 and BAL B-1b cells were quantified by flow cytometry (H) and lung mRNA by PCR (I,J). n=4-6 per group. *=p<0.05; **=p<0.01; ***=p<0.001.

Importantly, we also noted that treatment with bleomycin led to a drop in pleural B-1b cells (Figure 4C) as well as an increase in BAL B-1b cells (Figure 4H). This led us to hypothesize that loss of Id3 may poise pleural B-1b cell to be particularly responsive to lung injury. Given our findings showing the regulation of B-1 CXCR4 expression during lung injury, we asked whether Id3 may regulate CXCR4 expression. Indeed, loss of Id3 lead to increased surface CXCR4 expression across multiple B cell populations (Figure 5). Id3 regulates gene expression through interaction with E-proteins, transcription factors which bind E-boxes with the canonical CANNTG sequence. The CXCR4 promoter contains four such cis-regulatory elements. Therefore, we tested whether Id3 could directly regulate CXCR4 expression using the luciferase promoter reporter assay. The human CXCR4 promoter was cloned and transfected into the human Burkitt lymphoma cell line (BJAB). Transfection of E12 lead to induction of CXCR4 expression, while transfection of Id3 lead to a dose dependent inhibition of this induction. Taken together, these data establish Id3 as a novel regulator of CXCR4 expression and support the use of the Id3^BKO^ mouse as a model of augmented B-1b cell function and natural IgM.

**Figure 5.**
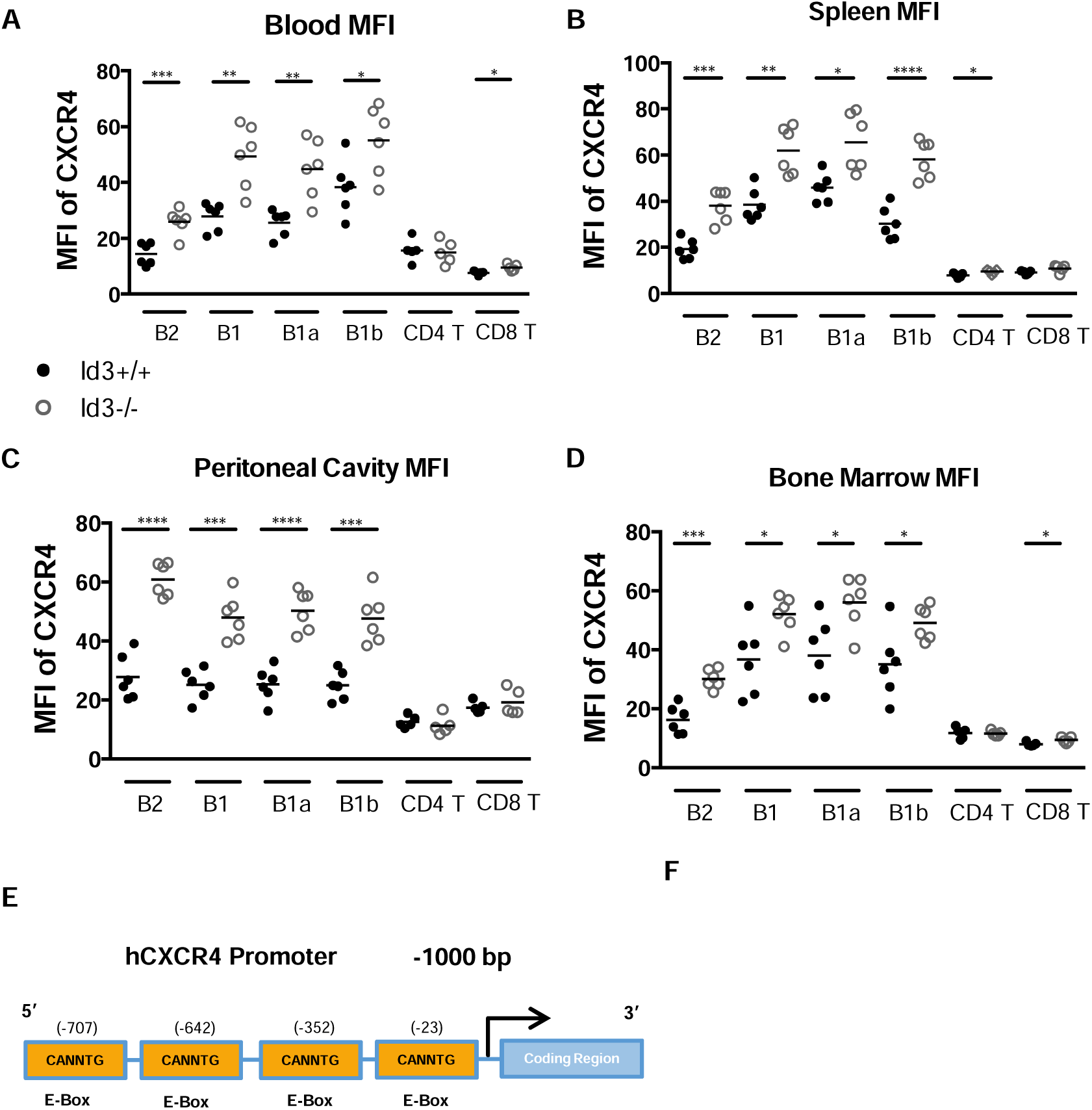
Id3 is a novel negative regulator of CXCR4 expression. (A-D) Flow cytometry of B and T cells from wild-type and *Id3^-/-^*mice showing increased CXCR4 surface expression in Id3-deficiency across multiple tissues. n=6 per group. (E) Illustration depicting consensus CANNTG (E-Box) HLH-transcription factor binding sites located within the first 1000 bases of the hCxCR4 promoter region. (F) BJAB cells were transfected with 1 μg of the CxCR4 promoter-luciferase reporter together with 2 µg of E12 and specified concentrations of Id3 and/or pcDNA3.1 empty vector. The presence of Id3 suppressed the promoter activity of CxCR4 in a dose-dependent manner. Luciferase activity is normalized to protein levels. Data are the result of 3 separate experiments of duplicate samples. Data represent mean ± SD; *=p<0.05; **=p<0.01; ***=p<0.001.

### B-1 cells and IgM inhibit lung injury and fibrosis

To test the effect of increased B-1b cells and natural IgM on fibrosis, we treated Id3^BKO^ mice with bleomycin and comprehensively assessed the development of lung fibrosis. Gene expression analysis showed that Col3a1 mRNA was reduced in Id3^BKO^ relative to WT mice (Figure 6A). MicroCT was performed to evaluate lung density throughout the lung (Figure 6B, representative images), and this showed reduced lung density in Id3^BKO^ mice, consistent with decreased fibrosis (Figure 6C). Finally, histologic analysis by Mason’s trichrome and fibrosis quantification by Modified Ashcroft also showed reduced fibrosis in Id3^BKO^ mice (Figure 6D-E). Collectively, these results show that increased B-1b cells and natural IgM can protect against the development of fibrosis.

**Figure 6.**
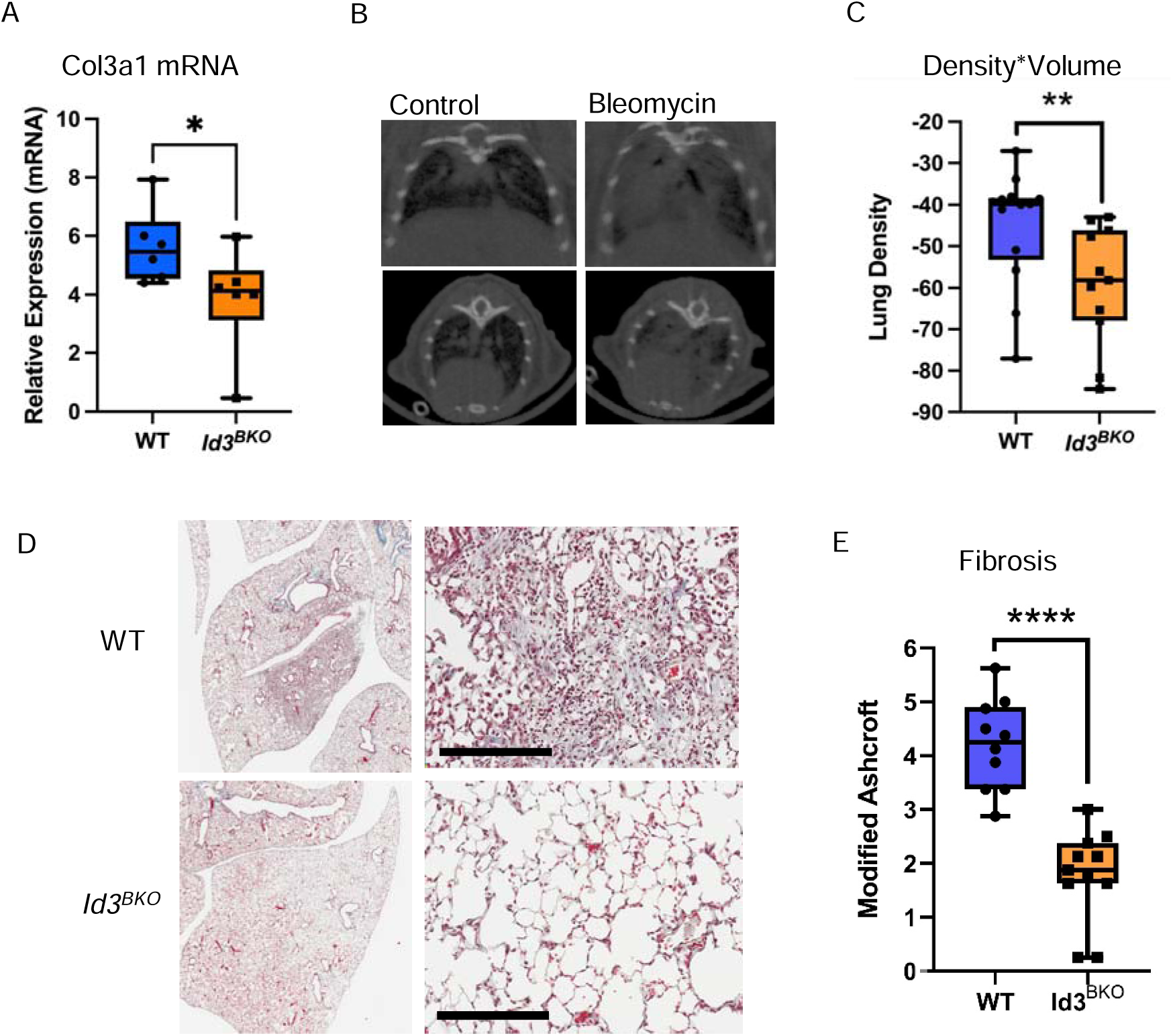
B cell-specific Id3-deficiency protects from bleomycin-induced lung fibrosis. *Id3^BKO^* mice and *WT* littermate controls were treated with bleomycin. At 21 days microCT scan was performed and density quantified (B,C). Lung RNA was harvested (A), and lung fibrosis was assessed histologically by Masson’s trichrome stain (D), quantified by modified Ashcroft score (E). *=p<0.05; **=p<0.01; ***=p<0.001. Scale bar = 200 μm.

In order to test the converse, IgM deficiency, we utilized *sIgM^-/-^*mice, which lack secreted IgM^36^. Treatment with bleomycin in *sIgM^-/-^* mice resulted in significantly increased weight loss and mortality (Figure 7) consistent with worsening lung injury. This could be partially rescued by delivery of E06 to the airway. Taken together with the data from Id3^BKO^ mice, these studies show that natural IgM, particularly oxidation-specific epitope IgM, plays a critical role in limiting lung injury and fibrosis.

**Figure 7.**
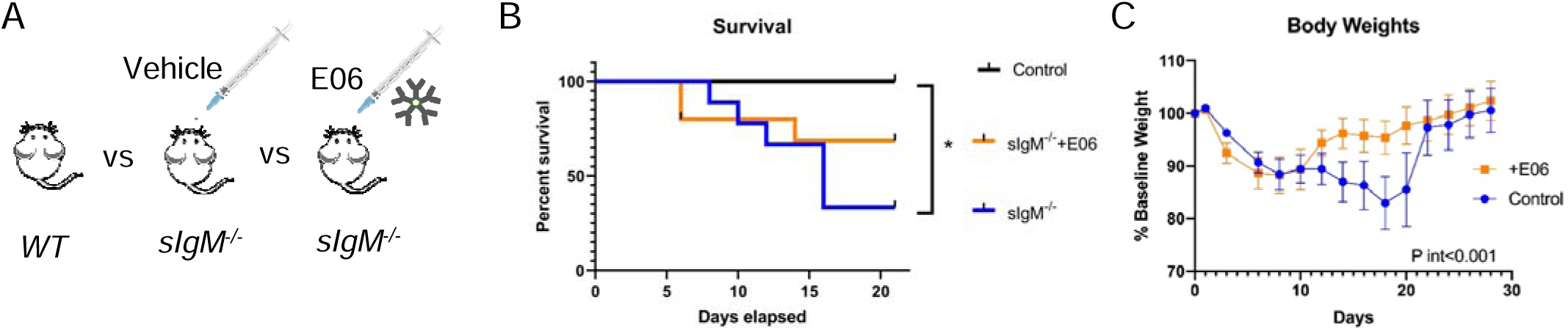
Delivery of the IgM^OSE^ E06 to the airway of IgM-deficient mice partially rescues lung injury. (A) Cartoon depicting the treatment groups. Survival (B) and weight (C) curves following bleomycin treatment. *=p<0.05.

We noted that the survival curves in the *sIgM^-/-^* mice showed a particular separation and increase in mortality after day 14 as compared to E06 treated mice. This timepoint is after the initial inflammatory response and considered to represent active fibroproliferation. Therefore, to probe potential mechanisms by which IgM may act to prevent fibrosis and promote healthy tissue repair, we treated *sIgM^-/-^*, *WT*, and Id3^BKO^ mice with and without bleomycin, and then harvested lungs at day 14 for bulk RNA sequencing. This allowed us to test the effect of a full spectrum of IgM levels on lung repair (Figure 8A). A heatmap of gene expression (Figure 8B) shows altered regulation of a wide range of genes. This is represented graphically by PCA analysis (Figure 8C). In this case groups are clearly segregated by bleomycin treatment, as well as sIgM deficiency. We then performed a GeneSet enrichment analysis of both experimental groups compared to WT mice, and we isolated those GeneSets which were significantly differentially regulated in response to bleomycin in opposite directions based on IgM. That is, only those genes that were up in *sIgM^-/-^*and down in Id3^BKO^, or down in *sIgM^-/-^* and up in Id3^BKO^ relative to WT are listed. Interestingly, the GeneSets which were inhibited by IgM (increased in *sIgM^-/-^*, decreased in Id3^BKO^; above the line) were primarily extracellular matrix and collagen formation, consistent again with blockade of fibrosis by IgM. GeneSets which were promoted by IgM (decreased in *sIgM^-/-^*, increased in Id3^BKO^; below the line) are notable for complement and complement activation.

**Figure 8.**
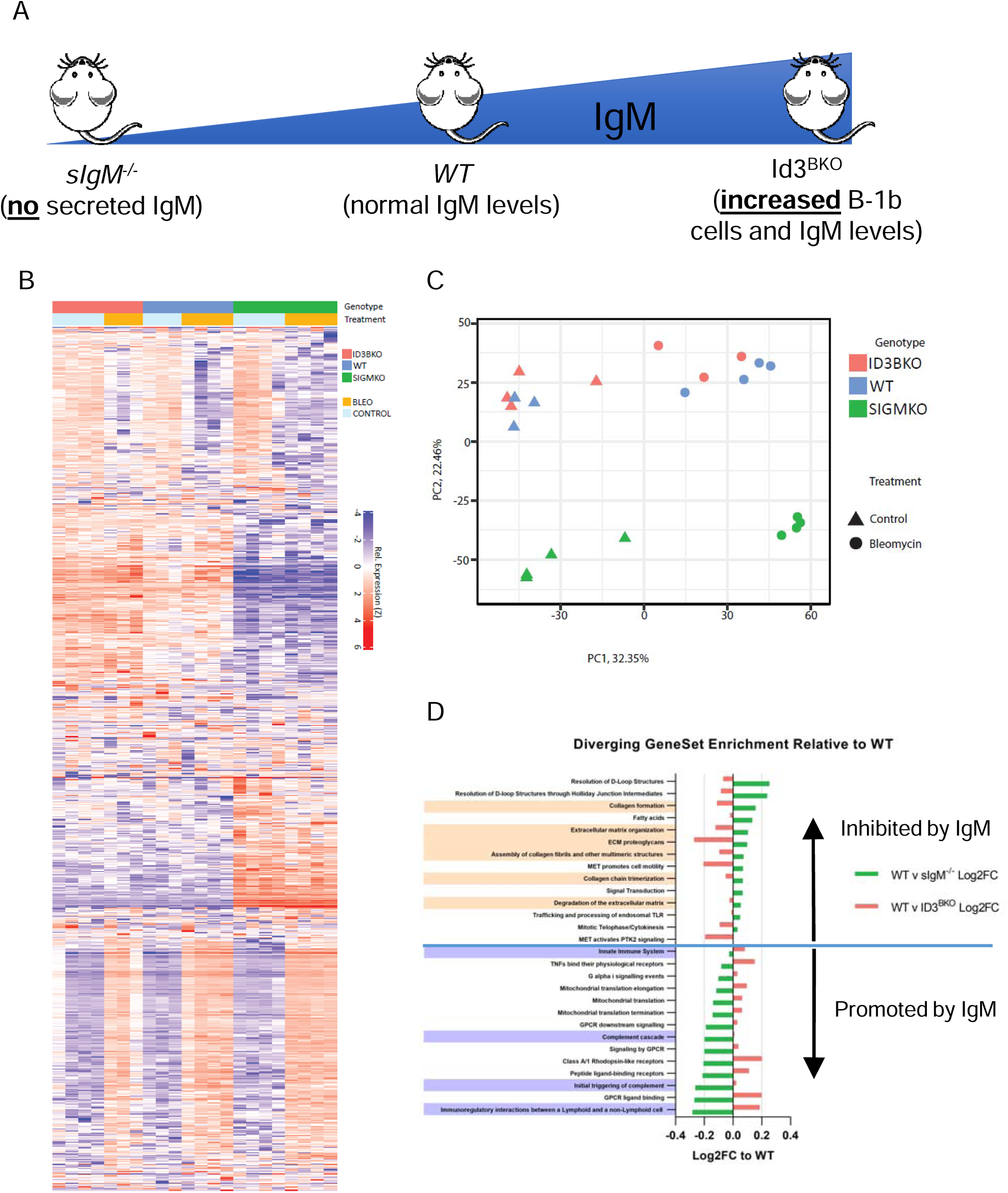
Lung RNAseq analysis across the spectrum of IgM gain- and loss-of-function. (A) Schematic showing the experimental design with three groups of mice: *WT*, *sIgM^-/-^*, and Id3^BKO^; and 2 conditions, control and bleomycin treated. (B) Heat map of gene expression across all groups. (C) Principal components analysis (PCA) of gene expression. (D) GeneSet enrichment analysis sorted by those inhibited by IgM (above the line) and promoted by IgM (below the line). n=3-4 mice per group.

**Figure 9.**
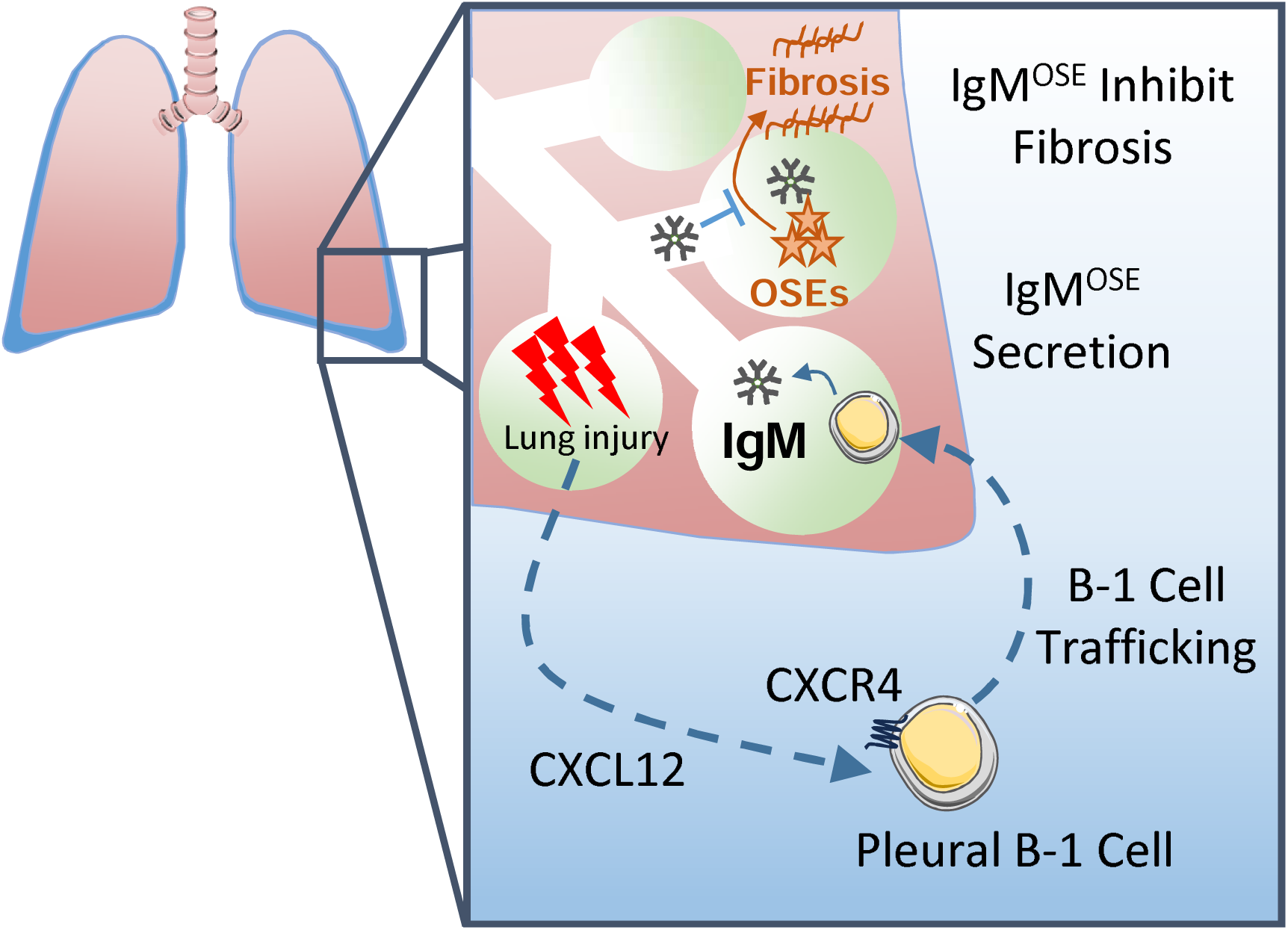
Proposed mechanism of the role of B-1 cells and IgM in lung injury and fibrosis. A lung injury leads to release of CXCR4 which signals resident pleural B-1 cells to migrate to the lung where they secrete IgM^OSE^. These antibodies block oxidation-specific epitopes such as those on damaged self to promote normal tissue repair and prevent the development of fibrosis.

## DISCUSSION

Dissecting the role of innate immunity in lung injury and repair is key to understanding the pathogenesis of lung fibrosis and designing new therapeutic interventions. Prior work has shown a clear role for DAMP signaling, in particular OSEs, in acute lung injury and pulmonary fibrosis. OSEs are generated during acute lung injury and can drive fibrosis^23, 37^. Here, we show that B-1 cells and IgM^OSE^, mediators of the innate immune response to tissue injury, play a key role in experimental lung injury and fibrosis. We show that B-1 cells are recruited to the injured lung where they secrete IgM and IgM^OSE^. We also show that this recruitment may involve the CXCR4-CXCL12 axis, as resident pleural and pulmonary B-1 cells upregulate surface expression of CXCR4 in response to injury.

To study the effect of B-1 cell recruitment to the injured lung, we utilized the Id3^BKO^ mouse as a viable model of increased pulmonary B-1 cells, and in so doing also identify Id3 as a novel transcriptional regulator of CXCR4 expression. This finding is relevant, not only for the role of B-1 cells in lung injury, but also more broadly as CXCR4 is a widely expressed chemokine receptor^38^ with potential roles in numerous cell-types and disease states, including HIV^39^, cancer^40^, and atherosclerosis^41^. In this model, we show that boosted levels of pulmonary B-1 cells and IgM are protective from fibrosis. Conversely, loss of secreted IgM worsens lung injury which can be partially rescued by airway delivery of IgM^OSE^. It is also possible that B-1 cells may have alternative roles in tissue remodeling which are independent of secreted IgM, such as paracrine cytokine secretion and antigen presentation, and these possibilities warrant further study.

Mechanistic studies into the role of secreted IgM in lung remodeling validated this protective role, showing an enrichment of extracellular matrix gene expression in the absence of IgM, and also point toward a role for complement as these related pathways were universally down-regulated in the absence of IgM. Terminal complement components have also been recently shown to be upregulated in BAL and plasma from patients with IPF^42^. What role this plays in disease pathogenesis remains to be determined. Our murine data suggests that activation of complement through IgM may be an important component of the normal repair response to tissue injury, and that loss of this axis leads to fibrosis. This may involve complement-mediated clearance of apoptotic cells, which IgM^OSE^ have been shown to bind^43^. This role of B-1 cells and IgM in tissue remodeling may extend outside of fibrosis. Recent data from human embryonic and fetal tissue identified higher levels of putative B1 cells in early embryonic development, suggesting they may also have a role in early organogenesis and tissue remodeling^44, 45^.

In summary, we show that B-1 cells and secreted IgM play key roles in the response to lung tissue injury and repair, and that augmentation of this axis can protect from the development of fibrosis. Subsequent studies should examine the role of B1-like cells and IgM in human fibrotic lung disease such as IPF, and further test the mechanistic role of complement activation in fibrosis.

## METHODS

### Mice

All animal studies were approved by the University of Virginia Animal Care and Use committee. Mice were obtained from Jackson Laboratory or bread in-house. All experiments were conducted with C57BL/6 background mice, ages 8-20 weeks, with genetic manipulations as described. Both sexes were used for all experiments unless otherwise specified. B cell-specific Id3 knock out mice (*Id3^flox/flox^;CD19^cre/+^*) were generated by crossing CD19^cre/+^ mice with Id3^flox/flox^ mice as previously described^46^.

### Bleomycin Model

Mice were sedated and bleomycin (1-2.5 U/kg) was administered trans-orally similar to previously described^47^. Following anesthesia, the mouse was suspended by the upper incisors, the tongue was retracted and 50 µl of bleomycin solution was delivered to the posterior oropharynx. With the tongue in the retracted position, the nares were covered until the solution was aspirated into the lungs. Following bleomycin administration, the mice were weighed daily and scored clinically across five categories including body weight, appearance, activity, respiratory, and posture. Human euthanasia endpoint was defined as 20% weight loss, a total score of 8 (out of 14 possible points), or a maximum score in 2 categories.

### Flow Cytometry

Mouse tissues were harvested and processed into single-cell suspensions as previously described ^33, 46^. Bone marrow and spleen were mechanically dissociated and passed through 70 µm screens. Lungs underwent mechanical digestion as well as enzymatic digestion with Collagenase and DNase.

### Quantitative Real-time Polymerase Chain Reaction

RNA was isolated using the RNeasy Mini Kit (QIAGEN) as per the manufacturer’s instructions. RNA was quantified, cDNA was made from equivalent starting amounts for each sample and synthesized using the iScript cDNA Synthesis Kit (Bio-Rad). Total cDNA was diluted 1:10 in water and amplified by real-time polymerase chain reaction using a BioRad CFX-96 iCycler and SYBER Green Supermix. Samples with no reverse transcriptase were used as negative controls. For comparison of relative mRNA expression levels, results were normalized to housekeeping genes18S ribosomal RNA and cyclophilin using the 2ΔΔCq method.

### Fibrosis Quantification

Micro computed tomography (CT) images were acquired using a Bruker Albira Si scanner (Billerica, MA). This scanner uses a rotating digital flat panel detector with a 50 kVp / 1 mA X-ray cone-beam generator. Images were reconstructed using a 3D based Filtered BackProjection algorithm. The resulting spatial resolution was 85 mm. Images were analyzed using AMIDE software. Lung fields were selected manually in axial cross-sections throughout the entire lung field and density was measured. For histologic analysis, mouse lungs were embedded in paraffin, sectioned into 10 µm, and stained with Masson’s trichrome. Sections were scored by Modified Ashcroft^48^ by two independent histologists who were blinded to the treatment group. A minimum of 10 fields per section at 20x magnification and 5 sections per mouse were scored and averaged.

### Cells

The human Burkitt lymphoma cell line (BJAB) was cultured in Dulbecco’s modified Eagle medium (DMEM,GIBCO) supplemented with 10% heat-inactivated fetal calf serum, 2 mM glutamine, and 100 U mL^-1^ penicillin and 100 mg mL^-1^ streptomycin at 37°C in an atmosphere of 5% C02/95% air.

### Luciferase Reporter Assay

BJAB cells were washed with serum free media and (2×10^6^ cells) were transfected by electroporation with 1 μg of the 935 bp h*CxCR4*-promoter-luciferase reporter (SwitchGear Genomics, Carlsbad CA) together with 2 µg total hE12 in pcDNA3.1 and increasing concentrations of hId3 in pcDNA3.1 and/or pcDNA3.1 empty vector. Electroporation was performed with a BioRad MXcell GenePulser (280 V, 950 µF, using the exponential waveform). 5 x 10^4^ transfected cells were plated in 6-well plates at 37C for 48 hrs. Cells were collected and washed in PBS, lysed, and assayed for luciferase activity. Protein concentrations from individual samples were quantified using the Pierce BCA Protein Assay Kit (Thermo Fisher Scientific) and luciferase values were normalized to protein.

### RNA sequencing

Cells isolated from murine lung were sent to Azenta Life Sciences (formerly GeneWiz), South Plainfield, NJ, USA for library preparation and sequencing. Extraction and Illumina library prep were performed with rRNA depletion. All 22 samples were sequenced across two lanes of the Illumina HiSeq sequencer to generate 2×150bp reads at a total read depth of 350 million paired-end reads per lane. Following sequencing, fastq files were processed to remove low-quality reads (Phred quality score < 20) and the presence of any adapter sequences using TrimGalore (*Babraham Bioinformatics - Trim Galore!*). The resulting quality of each sample was independently evaluated using FastQC^49^. Files that passed QC were further processed to obtain gene expression counts following a previously defined protocol by Pertea et al^50^. Briefly, reads were aligned to the mouse genome (UCSC mm10) using the HISAT2 aligner^51^ and transcripts assembled with StringTie^52^. The resulting StringTie output was used to produce read tables for further analysis.

### RNA Sequencing Analysis

The iDEP (integrated Differential Expression and Pathway analysis) platform (v1.1) was used for analyses^53^. Briefly, data were loaded and preprocessed using default parameters: 0.5 count-per-million (CPM) cutoff and transformed for clustering/PCA using EdgeR log2(CPM+4), with missing values imputed via median. DESeq2 was used for identification of differentially expressed genes (DEG) using an FDR of 0.1 and a min fold-change of 1.2^54^. GeneSet Enrichment Analysis (GSEA) was run using default parameters with the Reactome^55^ GeneSet library. Normalized Enrichment Scores (NES) were generated for each genotype for each condition. Fold changes in NES between wild-type vs ID3BKO and wild-type vs SIGMKO were pooled. Fold changes, thusly centered around wild-type as 1x, were log2 transformed and wild-type vs SIGMKO scores multiplied by negative 1 to allow for enrichment scoring along a continuum. The Log2FC scores were then sorted by maximum absolute value difference between ID3BKO and SIGMKO vs wild-type. *abs((log2FC(wt vs ID3BKO)) – (−1(log2FC(wt vs SIGMKO)),* providing an ordered list of the most differentially expressed genes along the ID3BKO(IgMhi) – wild-type (IgMwt) – SIGMKO (IgMlo) axis.

### ELISPOT

Enzyme-linked immune absorbent spot (ELISPOT) assays were performed as previously described ^33^. Plates were coated with capture antigen. Freshly isolated murine single-cell suspensions were then added to the plates, incubated overnight, and developed the following day with anti-IgM secondary.

### ELISA

Total IgM or antigen-specific IgM were measured from mouse plasma or BAL supernatants similar to previously described^33^. For oxidation-specific antibodies, plates were coated with either malondialdehyde-modified LDL or AB1-2 (the anti-idiotypic antibody specific for E06), then developed with anti-IgM secondary. Malondialdehyde-modified LDL was generated by modifying human LDL (Kalen Biomedical, LLC).

### Statistical Analysis

Statistical analyses were performed using GraphPad Prism software.

## Author contributions

J.M.S., R.T.H., A.U., E.O., E.T.F., A.A.E.A., C.T., V.J., J.C.G., A.M, and M.D.B. carried out experiments and performed data collection and analysis. J.M.S., R.T.H., Y.M.S., A.K., and C.A.M designed the studies. J.M.S. and C.A.M. obtained funding. J.M.S. wrote the manuscript. All authors participated in manuscript review and editing.

## Acknowledgments

The authors would like to thank Michael Solga from the UVA Flow Cytometry Core for technical assistance with flow cytometry, as well as Paul Soumen and Suart Berr from the Molecular Imaging Core.

---

<u>Supplemental Figure 1. Gating strategy for pulmonary B-1 cells</u>. Representative flow cytometry biaxial plots showing identification of B-1 cells as live singlets, CD45^+^; CD19^+^,CD3^-^; B220^mid-low^, IgM^high^. B-1a are CD5^+^, B-1b are CD5^-^.

