## Supplemental Figure 1 for "A protective role for B-1 cells and oxidation-specific epitope IgM in lung fibrosis"

### Slide 1
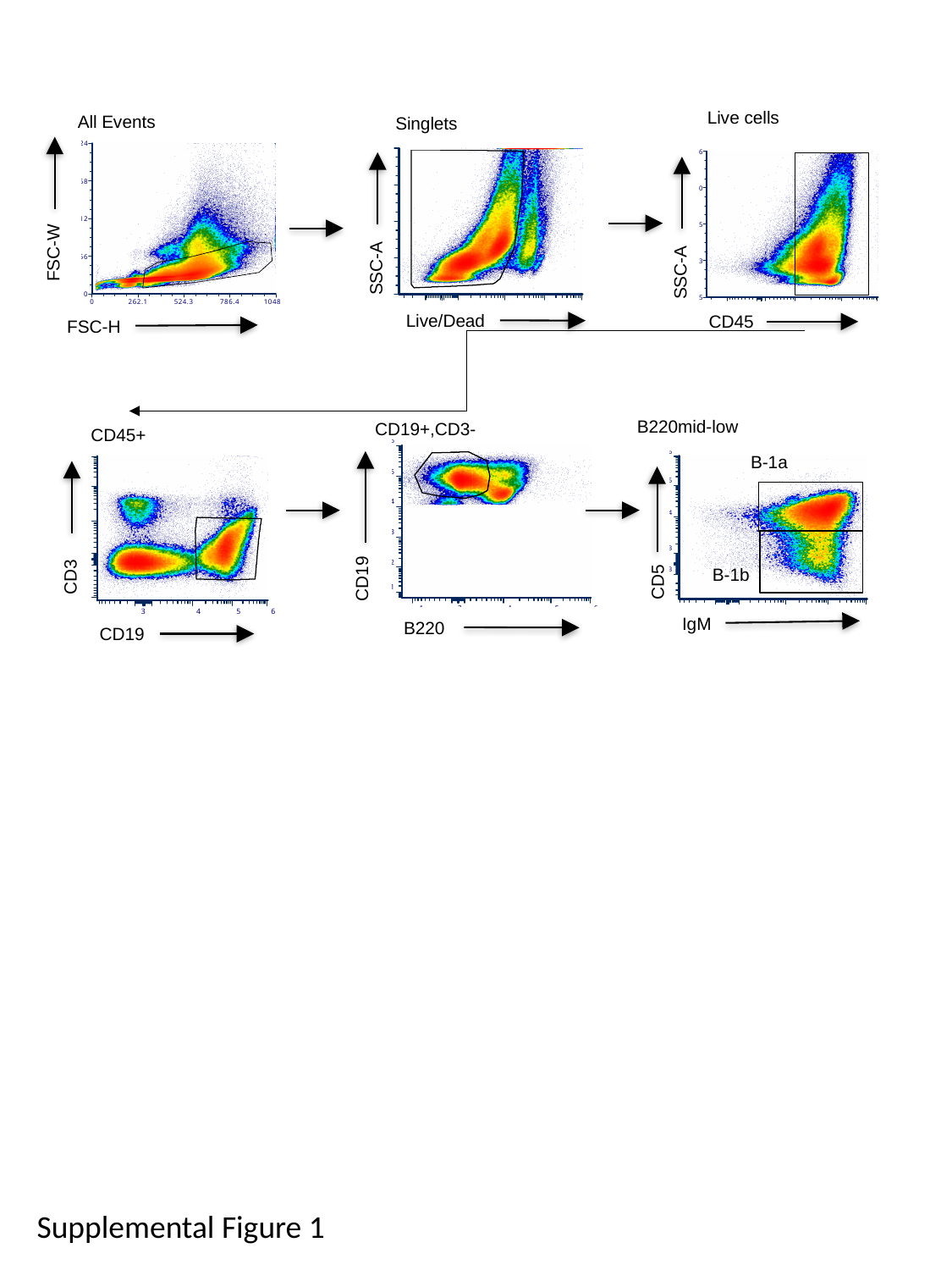

Live cells
SSC-A
CD45
All Events
FSC-W
FSC-H
Singlets
SSC-A
Live/Dead
B220mid-low
B-1a
CD5
B-1b
IgM
CD19+,CD3-
CD19
B220
CD45+
CD3
CD19
Supplemental Figure 1
